# Unveiling Distinct Neuroimmune Responses in Mouse Models of Cervical Spinal Cord Injury: Hemisection versus Hemicontusion

**DOI:** 10.1101/2025.08.02.668124

**Authors:** Wei Chen, Lucille Adam, Michel-Flutot Pauline, Arnaud Mansart, Stéphane Vinit, Isabelle Vivodtzev

**Affiliations:** Sorbonne Université, Development, Adaptation and aging (Dev2A), Unité CNRS UMR8263/Inserm U1345, 7 quai Saint Bernard, 75005, PARIS, France; Université Paris-Saclay, UVSQ, Inserm U1173, Infection et Inflammation (2I), 78000, Versailles, France; Inserm, END-ICAP, Université Paris-Saclay, UVSQ, 78000, Versailles, France

**Keywords:** Keyword: Spinal cord injury (SCI), cervical hemisection, cervical hemicontusion, mouse model, neuroinflammation, immune cell response, Chondroitin sulfate proteoglycans (CSPG), Glial migration, Macrophage polarization, systemic inflammation

## Abstract

Traumatic cervical spinal cord injury (cSCI) causes severe neurological deficits and long-term disability. Preclinical models such as cervical 2 (C2) hemisection (C2HS), resulting in disrupted communication between the respiratory centers and the phrenic motoneurons (PMN) pool, have been used since decades to study respiratory dysfunction and neuroinflammation after cSCI. Recently, contusive injuries such as C3 hemi-contusion (C3HC) have been increasingly used, as they induce phrenic motoneuron damage and offer a more clinically relevant model of SCI. However, these two different models may engage distinct pathophysiological cascades, raising concerns about the generalizability of findings across injury paradigms. In this study, we compared neuroimmune responses following C2HS or C3HC in mice. Animals underwent C2HS or C3HC, and spinal cord segments (C1-C8) were collected seven days post-injury for immuno-histological analyses around the lesion level and flow cytometry analyses at the lesion level. We observed that C2HS preserved more neurons and exhibited elevated CD86 and F4/80 expression. These markers are typically expressed by activated microglia and are indicative of a response oriented toward phagocytic and reparative functions. This phenotype was associated with limited pro-inflammatory cell infiltration and normalized level of systemic IL-6 in this model. Conversely, C3HC induced more extensive tissue damage, heightened microglial activation, a trend toward increased astrocytic reactivity, and significantly elevated CSPG levels on the contralateral side. Moreover, a persistent NK cell, neutrophil, and CD43⁺ antigen-presenting cells infiltration, along with persistently high circulating IL-6 has been observed following C3HC. These findings demonstrate distinct neuroinflammatory signatures and repairing mechanisms between models, with C2HS promoting a microglia profile toward repair and C3HC leading to a prolonged and potentially harmful immune response. This study underscores, for the first time, how injury type shapes neuroimmune mechanisms, reinforcing the need for lesion-specific therapeutic strategies in cervical spinal cord injury.

**Highlight:**

- C2 hemisection and C3 hemi-contusion trigger distinct neuroimmune responses in mice.
- C2HS preserves ventral neurons and upregulates CD86 and F4/80, suggesting repair-oriented microglia.
- C3HC induces CSPG accumulation, dendritic-like cell infiltration and prolonged systemic inflammation.
- Injury model influences neuroimmune environment and regenerative potential after cervical SCI.

## 1. Introduction

Spinal cord injury (SCI) induces a complex neuroinflammatory response, evolving through acute, subacute, and chronic phases (Anjum et al., 2020; Fan et al., 2022). Although inflammation is a hallmark of secondary injury, its characteristics vary depending on the nature of the trauma. Contusive injuries, which are most prevalent in human SCI, initiate distinct mechanical and biochemical cascades compared to transection models (Nicaise et al., 2012; Fan et al., 2022; Li et al., 2025). While transection models remain widely used for their high reproducibility when studying specific phenomena and molecular pathways, their differential neuroinflammatory impact on spinal microenvironment remains cryptic. Since inflammation influences both neural degeneration and repair, understanding these differences is essential for developing targeted therapeutic strategies (Donnelly and Popovich, 2008; Awad et al., 2013; Clifford et al., 2023).

This distinction is particularly relevant for studies of respiratory dysfunction, which have traditionally relied on the C2HS model due to its proximity to the phrenic motor circuits controlling diaphragm function (Vinit et al., 2006; Michel-Flutot et al., 2021). While this model has provided valuable mechanistic insights, its translational relevance has been questioned given that contusive or compressive injuries are more common in clinical SCI (Bajjig et al., 2022; Gayen et al., 2023). As a result, contusion models have gained increasing adoption due to their ability to better replicate the biomechanical forces and pathophysiology observed in human SCI. However, differences in neuroinflammatory profiles between these models are not yet well defined / or the extent to which these models diverge in their neuroinflammatory responses remains incompletely understood, raising concerns about the generalizability of findings across injury paradigms.

A fundamental question remains whether these distinct injury mechanisms, mechanical disruption in hemisection versus tissue damage in contusion, elicit divergent inflammatory responses, due to their importance in mediating subsequent repair processes. Of note, microglia, astrocytes, and infiltrating immune cells orchestrate the post-injury inflammatory cascade, yet their activation states, cytokine profiles, and spatial-temporal dynamics may differ between models (Perez et al., 2021; Dong et al., 2023). In addition, astrocytes and microglia are among the earliest responders after SCI, contributing both to neurotoxicity and tissue repair (Karimi-Abdolrezaee and Billakanti, 2012; Brockie et al., 2024). Following SCI, though astrocyte reactivity leads to the formation of a glial scar, involved in axonal regrowth inhibition(Ohtake et al., 2016a; Bradbury and Burnside, 2019; Yang et al., 2020), they can display protective abilities by maintaining tissue integrity and promoting healing(Yu et al., 2021; Smejkalova et al., 2025). Reactive astrocytes are also known to increase their chondroitin sulfate proteoglycans (CSPGs) production, which form one of the major components of inhibitory perineuronal nets. These specialized extracellular matrix structures stabilize synaptic connections but limit neuronal plasticity and regeneration (Fawcett et al., 2019; Hosseini et al., 2024). These CSPGs can be visualized using Wisteria floribunda agglutinin (WFA), a lectin that binds specifically to N-acetylgalactosamine residues within CSPG structures, making it a widely used marker for perineuronal nets and inhibitory matrix components following CNS injury (Irvine and Kwok, 2018; Mukhamedshina et al., 2019). Microglia on their end, can exhibit a range of activation phenotypes simultaneously, including both pro-inflammatory states that contribute to secondary injury, and anti-inflammatory (or reparative) states that support tissue recovery.

In addition to resident glia, the infiltration of peripheral immune cells is a hallmark of secondary injury. Neutrophils are quickly recruited at the lesion site, exacerbating tissue damage via protease and reactive oxygen species release (Rawat and Shrivastava, 2022). Natural killer (NK) cells and monocytes-derived cells are subsequently recruited and involved in modulating the immune response and influence both neurodegeneration and repair (Bowes and Yip, 2014; Garofalo et al., 2020). Of note, surface markers such as CD68, CD86, F4/80, and CD206 enable characterization of these immune populations, distinguishing pro-inflammatory from reparative phenotypes (Gao et al., 2022; Lee et al., 2023). Hence, elucidating the recruitment, distribution, and activation profiles of these immune populations is critical for understanding injury-specific immune dynamics and for developing targeted immunomodulatory therapies.

In this study, we directly compare neuroinflammatory responses in two clinically relevant cervical SCI models: the historically used C2 hemisection model and a more recently developed C3 hemi-contusion model. By analyzing innate immune activation (microglia, astrocytes, macrophages) and quantifying key inflammatory mediators both at and below the lesion epicenter, we defined model-specific inflammatory patterns and we shed light on distinctive inflammatory mechanisms occurring post-SCI in two of the most used SCI preclinical models.

## 2. Material and methods

### 2.1 Animals

Adult male Swiss mice (30-40 g; 6 weeks) were used for in vivo studies. Animals were dually housed in ventilated cages in a state-of-the-art animal care facility (2CARE animal facility, accreditation A78-322-3, France) under a 12 h light/dark cycle with ad libitum access to food and water. All procedures were performed in accordance with the European Communities Council Directive (Surgical Procedures), approved by the Ethics Committee Charles Darwin CEEACD/N 5 (Project authorization APAFIS No. 201901301500576 and 2021080910063249).

### 2.2 Study design

A total of 36 adult male mice were used to compare neuroinflammatory responses between two cervical spinal cord injury (SCI) models: C2 hemisection (C2HS) and C3 hemicontusion (C3HC). Two complementary approaches were applied. Immunofluorescence staining was performed on fixed spinal cord tissue collected from segments at and below the lesion site to evaluate glial activation and extracellular matrix changes. This cohort included mice that received laminectomy only as controls (n = 2), C2HS (n = 6), and C3HC (n = 11). In parallel, flow cytometry was conducted on fresh tissue isolated from the lesion epicenter to quantify resident and infiltrating immune cell populations. The flow cytometry cohort consisted of control mice (n = 4), C2HS (n = 7), and C3HC (n = 6). This combined experimental design allowed for a comprehensive evaluation of injury-specific immune responses at the cervical level.

### 2.3 C2HS and C3HC models

Animals were anesthetized with isoflurane (5% in 100% O2) in a closed chamber, then maintained throughout the surgical procedure with isoflurane (2.5% in 100% O2). After skin and muscles were retracted, laminectomy and durotomy were performed at the C2 level for C2HS and at C3/C4 for C3HC. For the hemisection model, the spinal cord was then sectioned unilaterally (left side) with microscissors. To ensure the section of potentially remaining fibers, a microscalpel was used immediately after microscissors, as described previously(Fayssoil et al., 2021).For the contusion model, a precision Impactor Device (RWD life science; 68,099 II) with a 1.5 mm tip impactor was used to perform the hemicontusion as previously described (Bajjig et al., 2022). The impactor parameters had a mean depth of 2.6 ± 0.2 mm, a speed of 1.0 ± 0.2 m/s and a dwell time of 0.60 ± 0.01 s (mean ± SEM) for the injury After section, sutures were used to close the wounds and skin.

Post-operative care was standardized for both lesion models. Animals were placed on a heated pad to recover, then returned to their home cage with *ad libitum* access to soft food and water (or jellified water for the 1^st^ post-operative day). Solid food and water were also provided on the top grid of the cage. Subcutaneous fluids were administered as needed to prevent dehydration on the first few post-operative days. Body weight and food intake were monitored daily to ensure the best environment possible for recovery. All mice received the same standardized antibiotic and analgesic regimen: subcutaneous injections of sulfadoxine (Borgal, 0.2 mL per mouse; 0.1 mL diluted in 1 mL NaCl) and buprenorphine (Buprécare, 0.5 mL per mouse; 0.05 mL diluted in 1 mL NaCl) for infection prevention and analgesia respectively.

### 2.4 Immunofluorescence staining and analysis

Animals were euthanized by intracardiac injection of pentobarbital (EXAGON, Axience), intracardially perfused with heparinized 0.9% saline, followed 4% paraformaldehyde. Spinal cords were dissected and post-fixed in 4% PFA overnight at 4 °C for 24 h, rinsed with PBS and cryoprotected in 30% (in 0.9% NaCl) sucrose for 24 h. Transversal free-floating slices (30 µm) were obtained using a Thermo Fisher NX70 cryostat. Spinal cord slices were stored in a cryoprotectant solution (30% sucrose, 30% ethylene glycol, 1% polyvinylpyrrolidone in 1X PBS) at -20 °C. 30 µm thickness slices were collected from C1 to C8. For immunofluorescence experiments, free-floating transverse sections of the C1- C8 spinal cord were washed and placed in blocking solution (Normal donkey serum (NDS) 5% with Triton X100 0.2% in PBS 1X) for 30 min. After blocking, sections were incubated with primary antibodies overnight on an orbital shaker at 4 °C and followed by a 2 h incubation in secondary antibodies solution at room temperature. Primary and secondary antibodies used in this study are detailed in Supplementary Table 1. Slices were then incubated with NeuroTrace™530/615 (N21482, Invitrogen, Waltham, MA, USA), a neuronal marker (Nissl stain), for 10 min and then washed again 3 times with PBS 1X. For each staining, 6 individual animals per group were examined and images were captured with a Hamamatsu ORCA-R2 camera mounted on an Olympus IX83 P2ZF microscope or a 3dhistech panoramic slide scanner. Images were analyzed using ImageJ 1.53n software (NIH, USA).

### 2.5 Flow cytometry staining/gating and analysis

Animals were euthanized by heart exsanguination with heparinized syringe under isoflurane anesthesia. After opening the blood circulation, systemic blood was flushed from the tissues by perfusing the left heart ventricle with 1X PBS-EDTA (2 mM). Immediately after perfusion, spinal cord section centered around the lesion epicenter was dissected out (around 20mg per tissue segment) and placed in collagenase D (2.5 mg/mL) prepared in RPMI 1640 medium (Roswell Park Memorial Institute, catalog no. 21875-034, Thermo Fisher Scientific), minced with microscissors and digested for 20 min at 37°C. Tissue was mechanically dissociated and passed through a 100 μm nylon mesh to obtain a single-cell suspension. Cells were then washed with RPMI and resuspended in PBS 1X for antibody labeling. Viability staining was performed with Fixable Viability Dye eFluor™ 780 (1/2000, 65-0865-18, Invitrogen, Waltham, MA, USA) Cells were then incubated with a cocktail of fluorescent antibodies for 30 minutes at 4°C in PBS 2% fetal calf serum (FCS) (antibodies are detailed in Supplementary Table 2- Panel 1 and Panel 2). Cells were washed with PBS FCS 2% and passed through a 40um cell mesh before acquisition with a BD LSRFortessa™ III (BD Life Sciences, Franklin Lakes, NJ, USA). Data was analyzed with FlowJo™ v10.8.1 Software (BD Life Sciences, Franklin Lakes, NJ, USA) software.

### 2.6 Cytokine and chemokine expression analysis

Blood was collected on anesthetized animal via intracardiac route just before euthanasia (7 days post injury). Plasma was prepared from blood obtained during euthanasia and stored at -80°C until analysis using multiplex (macrophage/microglia LEGENDplex™ panel, BioLegend®) according to manufacturer’s instructions with slight modification: all reagent and sample volumes were divided by two except washing volumes. Analysis was performed using LEGENDplex™ Data Analysis Software Suite.

### 2.7 Statistical analysis

Numerical values are reported as mean ± standard deviation (SD). Datasets were tested for normality using Shapiro–Wilk test. One-way ANOVA tests for more than two groups when the data were normally distributed was performed. Mann-Whitney tests were performed for two-sample comparisons or Kruskal-Wallis tests for more than two samples if the data were not normally distributed. Normal distribution was assumed if sample size was sub-threshold for normality testing. Statistical analyses were performed with GraphPad Prism v10 software and difference were considered significant * at p < 0.05, ** at p < 0.01 and *** at p < 0.001. All statistics are stated in the figure legends and correction for multiple comparisons performed where appropriate.

## 3. Results

### 3.1 Neuroinflammatory and perineuronal environment below the lesion level

Due to the highly focal and anatomically restricted nature of the C2 hemisection, direct analysis at the lesion epicenter is technically challenging. Instead, we focused on tissue located below the lesion, which enables a consistent and anatomically comparable assessment across models, and allows better isolation of the effects related to injury level. This approach also provides valuable insight into secondary pathological changes and potential plasticity occurring in spared regions of the spinal cord. Immunofluorescence staining of microglia (Iba1) and astrocyte (GFAP) in the spinal cord below the lesion level is presented on figure 2. Quantification was performed by measuring the percentage of positive area occupied by Iba1 and GFAP immunoreactivity on each side of the spinal cord at ventral horn. We found a significant higher proportion of Iba1-positive area ipsilaterally to the injured side after C3HC compared to C2HS (36.6 ± 9.3 %, vs. 4.3 ± 0.7 %, p < 0.01). A trend toward increased in Iba1-positive area was also observed on the uninjured side in the C3HC model compared to C2HS: (20.3 ± 6.6 % vs. 5.0 ± 0.9 %, p = 0.08). Similarly, a significantly higher proportion of GFAP-positive area was found below the lesion in the injured side in the C3HC compared to the C2HS model (2.4 ± 0.6 % vs. 0.5 ± 0.1 %, p < 0.05), with no significant difference in the uninjured side below lesion level between injury models. These findings suggest a more pronounced microglia l and astrocytic response after C3HC.

**Figure 1.**
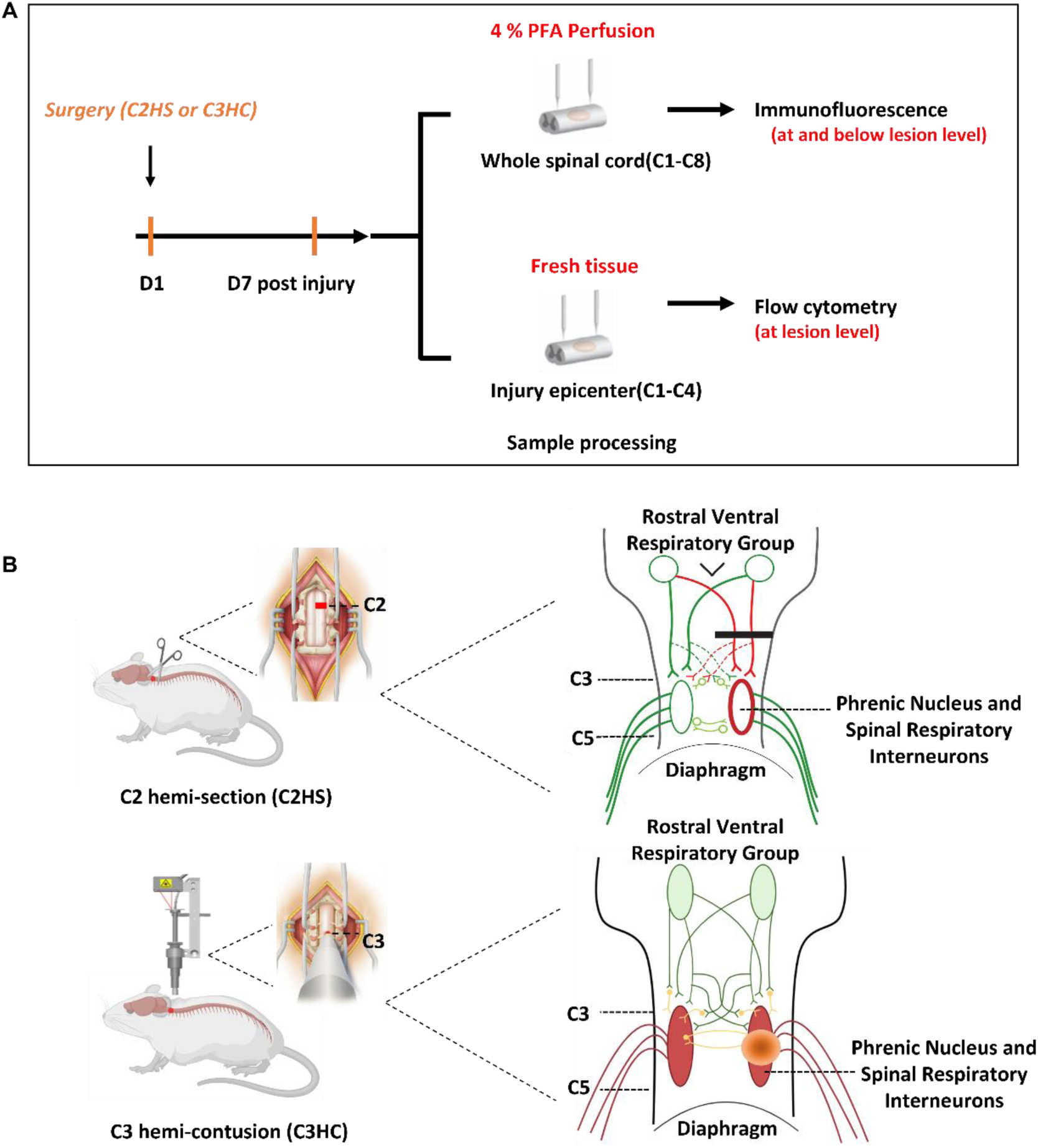
**A.** Experimental design of immune cell response after 7 days SCI study. **B.** Surgical exposure of cervical spinal cord injury models and Schematic representation of the two different procedures and their effect on the spinal network. Hemisection (C2HS), involving unilateral sectioning of the spinal cord at the C2 level, resulting in disconnection between respiratory centers and the phrenic motoneuron (PMN) pool at C3-C5; and the Hemicontusion (C3HC), involving unilateral contusive injury at the C3/C4 level, resulting in direct damage to PMNs and providing a clinically relevant SCI model.

**Figure 2.**
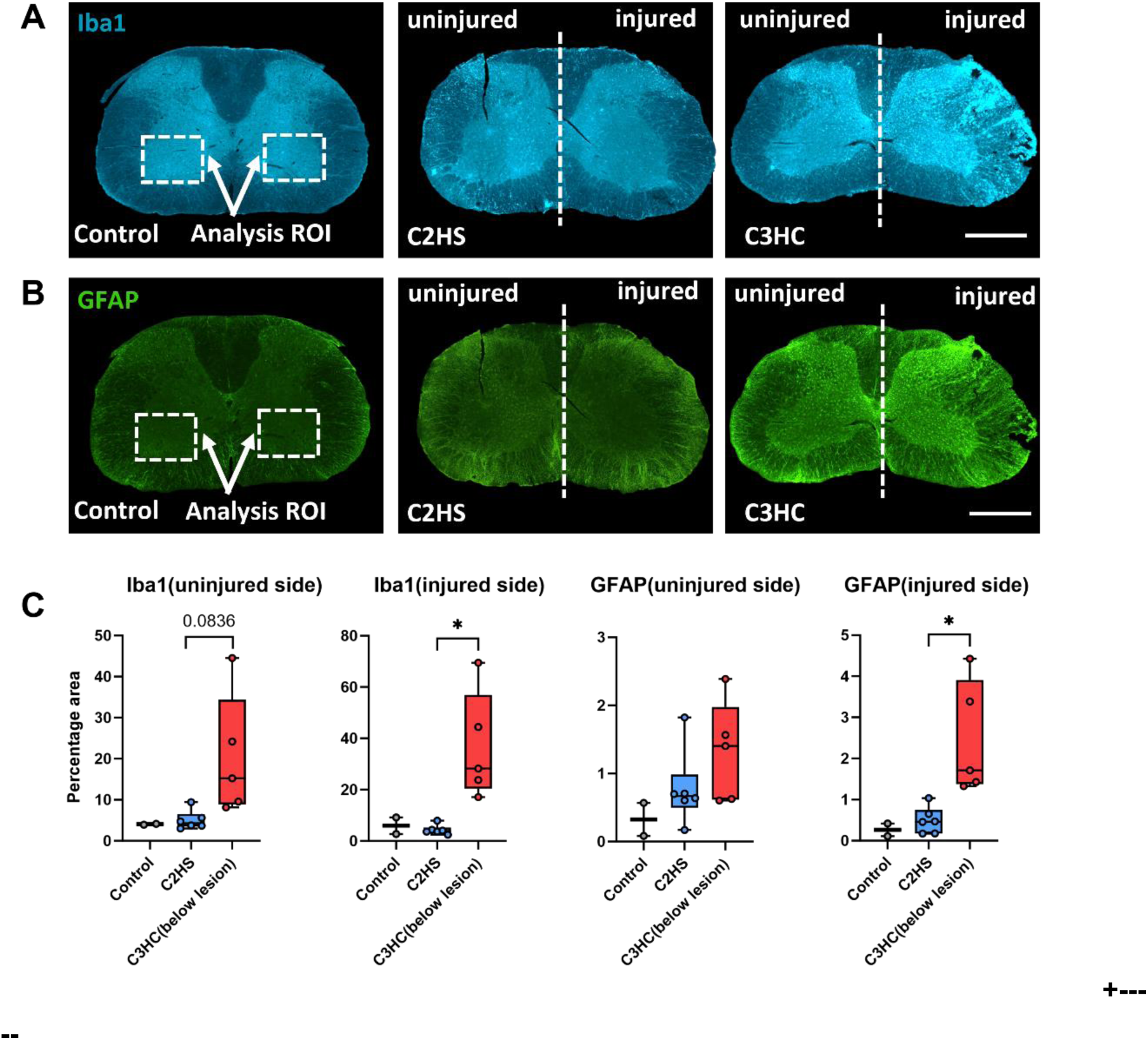
Microglia and astrocyte immunostaining at 7 days post-injury after C2HS vs.C3HC below the lesion level. **A.** Representative images of expression of ionized calcium-binding adaptor protein-1 (Iba1) (microglia, in blue) positive percentage area at ventral horn and **B.** expression of glial fibrillary acidic protein (GFAP) (Astrocyte, in green) positive percentage area at ventral horn for control, C2HS, C3HC group following 7 days post-injury below the lesion level (C3-C8 for C2HS; C5-C8 for C3HC). The white dashed boxes indicate the analysis region of interest (ROI) used for histogram quantification. In the C2HS images, a small incision is visible on the left dorsal horn, which was made during tissue processing to mark the uninjured side. **C.** Quantification of the percentage positive area of Iba1 and GFAP staining at ventral horn for uninjured and injured sides of C2HS and C3HC animals following 7 days post-injury below the lesion level. Scale bar: 500 µm. Statistical analysis was performed using the One-Way ANOVA and unpaired t test (*), * *p*<0.05, ** *p*<0.01, *** *p*<0.001. Scale bar: 500 µm.

Immunofluorescence staining of WFA in the spinal cord below the lesion level is presented on (Figure 3). A greater proportion of WFA-positive area was found below the lesion level after contusion on both sides compared to section (Injured: 34.8 ± 6.2 % vs. 14.2 ± 3.4 %, and uninjured: 32.0 ± 5.5 %, vs.: 10.8 ± 3.0 %), p < 0.05). However, the total number of neurons was significantly reduced after C3HC compared to C2HS (272 ± 11vs. 324 ± 6, p<0.01). As a result, the proportion of WFA-positive perineuronal structures were significantly different from control but did not differ significantly between the two groups. Additional results from the lesion epicenter, including a higher percentage area of Iba1- positive microglia, greater GFAP immunoreactivity, and less neuronal survival in the contusion model, are presented in Figures S2 and S3.

**Figure 3.**
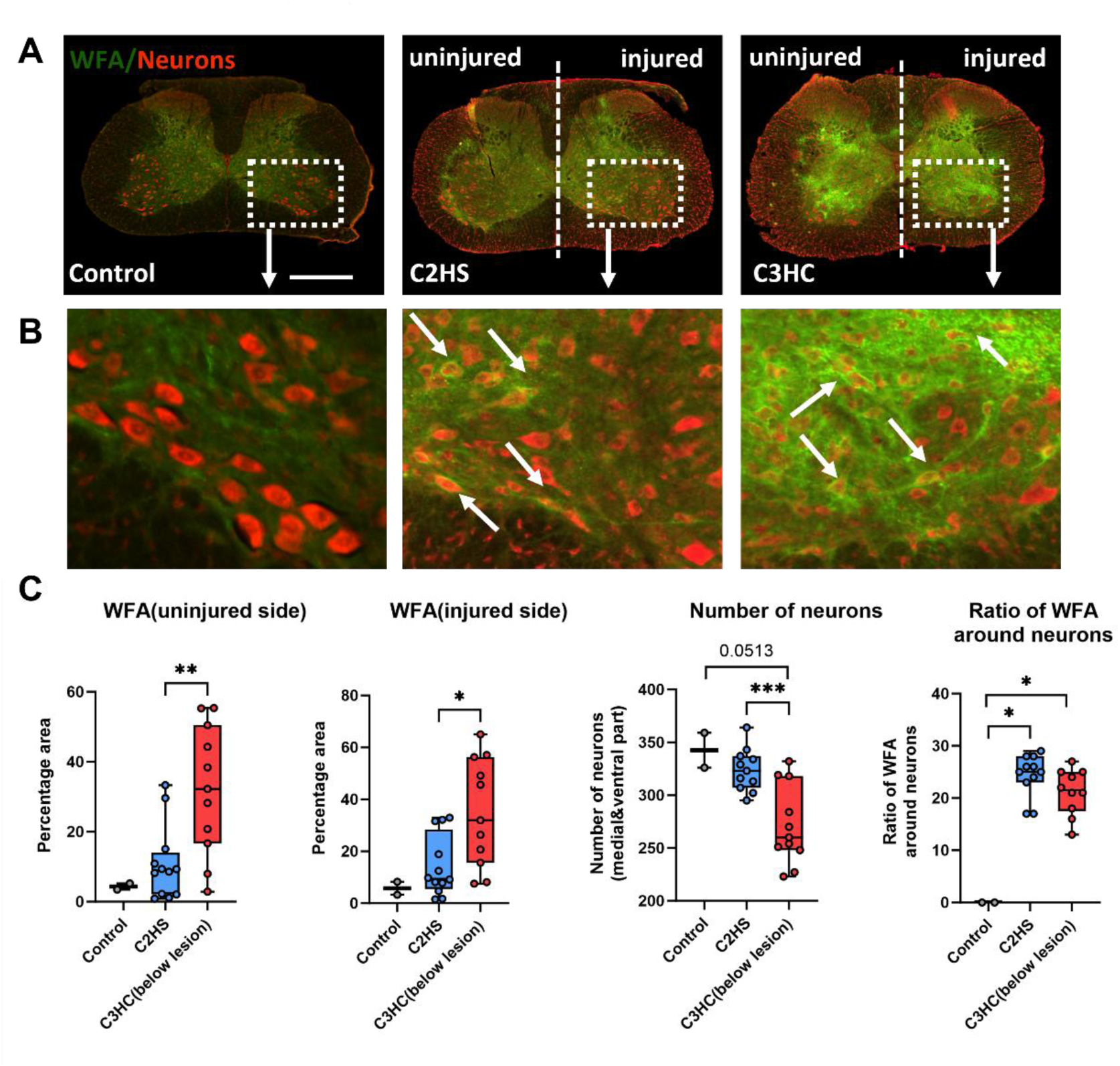
WFA staining and ventral neuron changes at 7 days post-injury in C2HS and C3HC cervical spinal cord injury models below the lesion level. **A.** Representative images of WFA staining around ventral neurons in control, C2HS, and C3HC groups at 7 days post-injury below the lesion level. **B.** Higher magnification images (10×) highlighting neurons surrounded by WFA-positive structures. **C.** Quantification of the percentage area occupied by WFA staining, number of ventral neurons, and proportion of neurons associated with WFA for uninjured and injured sides of C2HS and C3HC animals following 7 days post-injury below the lesion level. Scale bar: 500 µm. (*), * *p*<0.05, ** *p*<0.01, *** *p*<0.001.

### 3.3 Differences of cell population at the lesion level

We then characterized immune cell populations at lesion site by flow cytometry. Neutrophils, natural killer (NK) cells, microglia , and CD43⁺ cells were identified. CD43+ cells were further subdivided into subsets based on F4/80 and MHC-II expression to defined infiltrated antigen-presenting cells (APCs; CD43+ F4/80+ MHCII+), CD11b+ dendritic cell (DC)-like cells (CD43+ F4/80- MHCII+ CD11b+) and CD8+ DC-like cells (CD43+ F4/80- MHCII+ CD8+) (Supplementary Figure 1). Overall, these immune populations were increased in both injury models compared to controls, with a greater magnitude of infiltration observed at the contusion model compared to section model. (Figure 4). In the C3HC model, significantly higher numbers of Neutrophils were observed (HS: 73.4 ± 10.1 cells/mg vs HC: 125.7 ± 19.2 cells/mg, p < 0.05), as well as increased NK cell infiltration (HS: 9.3 ± 1.9 cells/mg vs. HC: 16 ± 1.5 cells/mg, p < 0.05) compared to the C2HS group. A significant elevation of Infiltrated APC (HS: 34 ± 7.9 cells/mg vs. HC: 88.8 ± 15.1 cells/mg, p < 0.05) was also noted in the C3HC model compared to the C2HS group. Among these, the CD11b⁺ DC-like subset was significantly increased (HS: 23.3 ± 4.6 cells/mg vs. HC: 49.9 ± 12.1 cells/mg, p = 0.05) and a trend toward increased numbers of CD8+ DC-like cells was also observed (HS: 9.0 ± 4.4 cells/mg vs. HC: 13.6 ± 6.9 cells/mg, n.s.), although the difference was not statistically significant. In contrast, at the lesion level, we found the number of Microglia did not differ significantly between groups.

**Figure 4.**
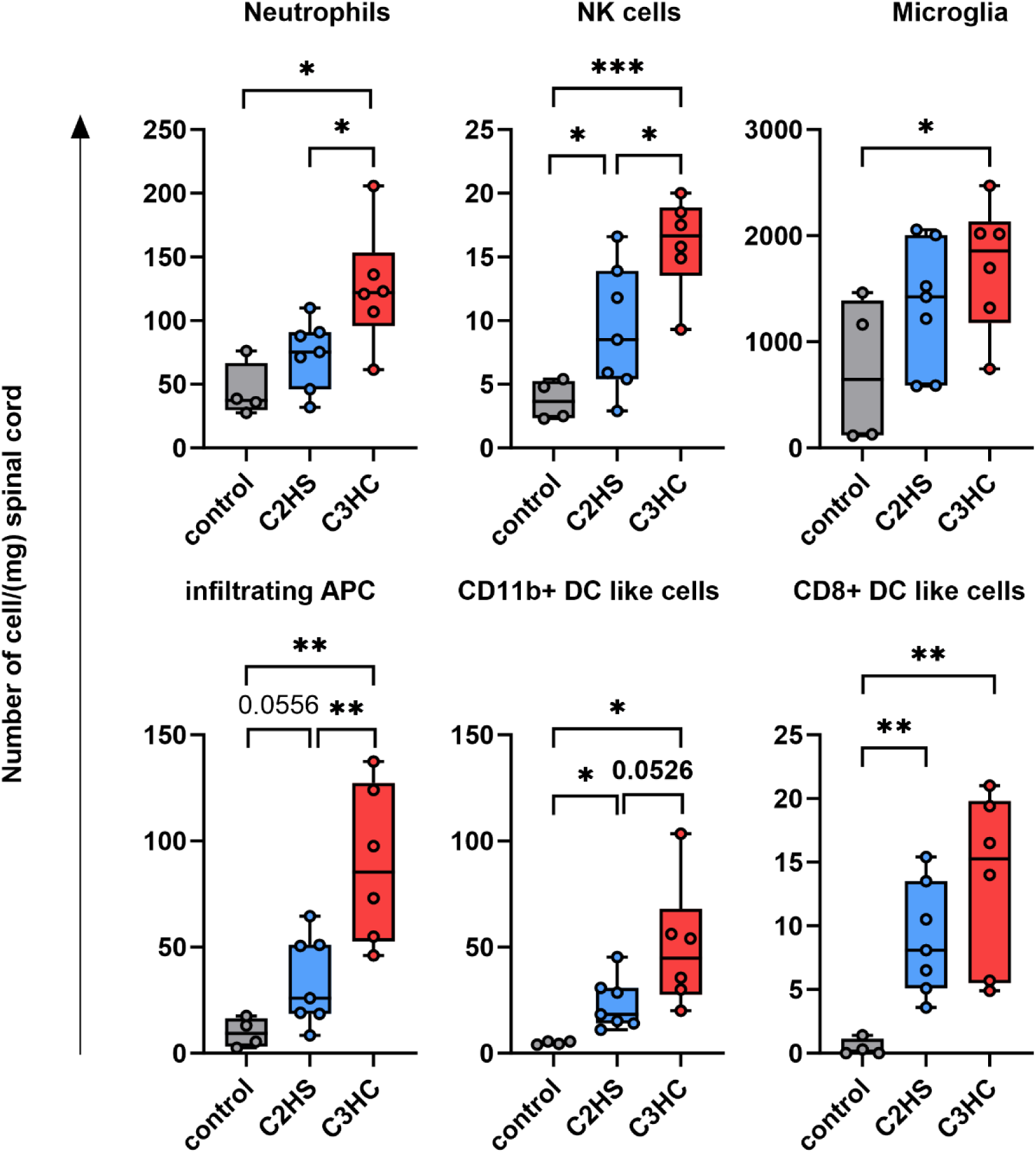
Characterization of the immune response at the spinal cord lesion level 7 days after injury. Histograms (Min to max, Box and Whiskers) showing the number of cells (Neutrophils, NK cells, Microglia, infiltrating cells, CD11b+ APC-like cells, and CD8+ APC-like cells) counted by mg of spinal cord tissue collected at the lesion site or at the equivalent site for Sham animals. Statistical analysis was performed using the One-Way ANOVA and unpaired t test (*), * *p*<0.05, ** *p*<0.01, *** *p*<0.001.

**Figure 5:**
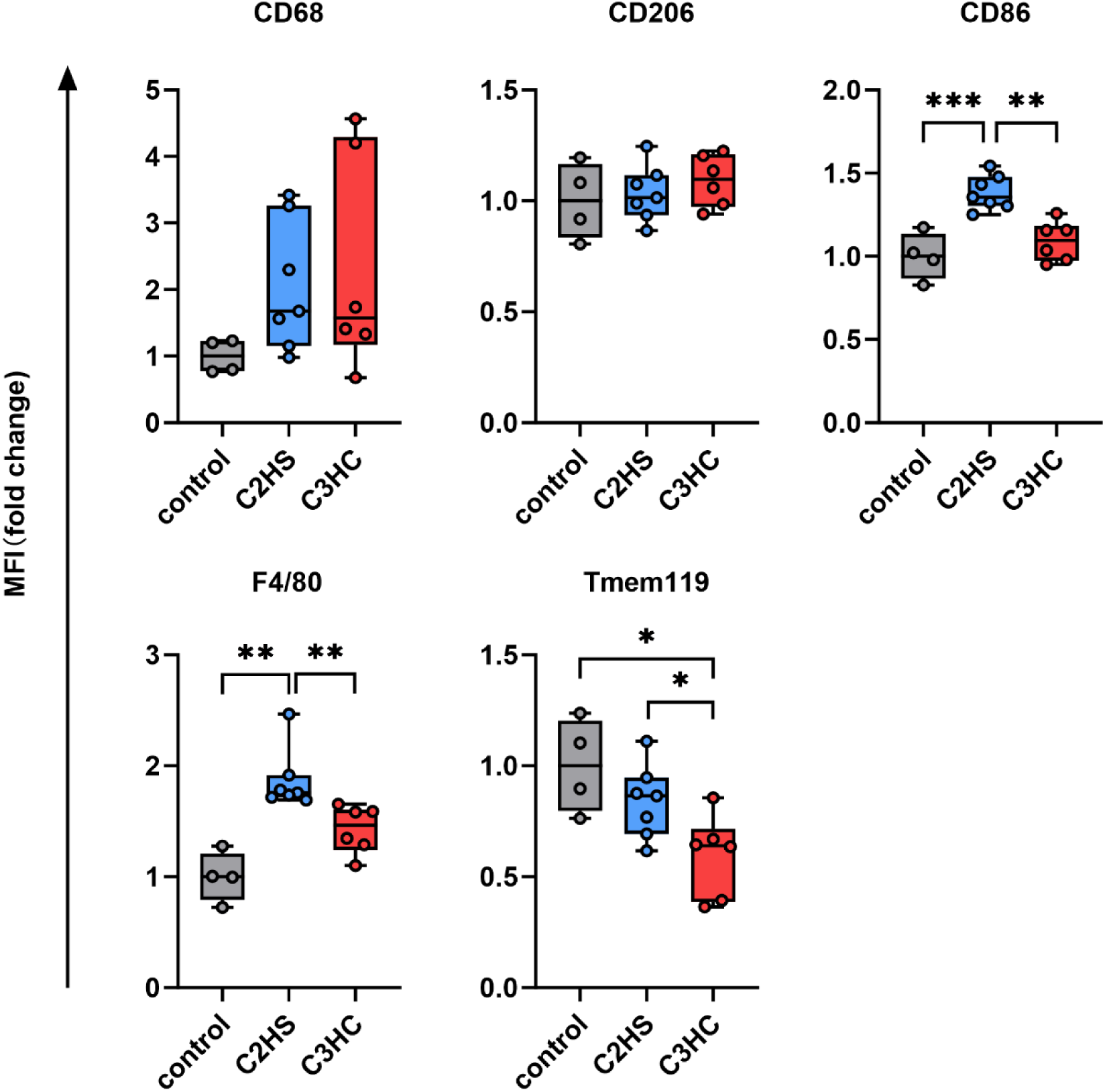
Microglial surface marker expression following cervical spinal cord injury for control, C2HS, and C3HC groups. Histogram (Box and Whiskers graphs (Min to max)) presenting the expression level of markers at Microglia cell surface. Data are expressed as fold change using Mean fluorescence intensity (MFI), normalized to the mean value of the control group. Statistical analysis was performed using the One-Way ANOVA and unpaired t test (*), * *p*<0.05, ** *p*<0.01, *** *p*<0.001.

**Figure 6.**
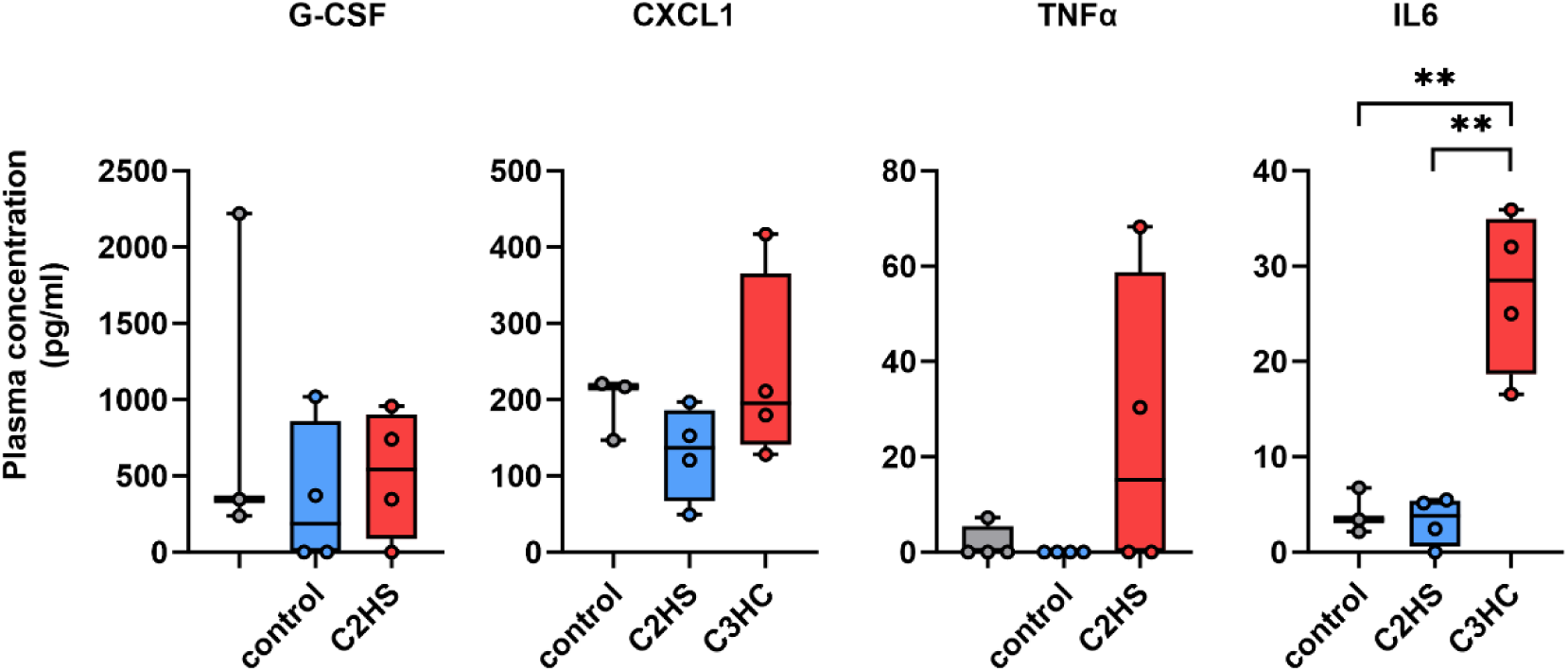
Circulating cytokine levels in control, C2HS, and C3HC mice at 7 days post-injury. Box and Whiskers graph (mini to max) representation of proinflammatory plasma level cytokines and chemokines at 7 days post C2HS, C3HC or control measured by LEGENDplex™. Statistical analysis was performed using the One-Way ANOVA and unpaired t test (*), * *p*<0.05, ** *p*<0.01, *** *p*<0.001.

Moreover, we studied Microglia activation state by measuring expression levels of surface marker mainly associated with inflammatory activation profile (M1 like) or homeostasis/ repair profile (M2 like). The mean fluorescence intensity (MFI) of CD68, a marker illustrating activation, trend to increase in Microglia at both injury model while CD206 MFI, a marker associated with anti-inflammatory macrophages remain comparable to control. Interestingly, CD86 and F4/80 was significantly higher in the C2HS group (HS: 1.4 ± 0.1 a.u. vs. HC: 1.1 ± 0.1 a.u., p < 0.01) and (HS: 1.9 ± 0.3 a.u. vs. HC: 1.4 ± 0.2 a.u., p < 0.01) respectively. The expression of these markers by macroglia have been associated with repair/phagocytosis function (Zhang et al., 2023). In addition, a reduction in Tmem119 expression, a marker of homeostatic Microglia, was more pronounced in the C3HC group (HS: 0.8 ± 0.2 a.u. vs. HC: 0.6 ± 0.2 a.u., p < 0.05).

Lastly, we assessed systemic inflammatory responses following C2HS and C3HC by measuring circulating levels of key cytokines: interleukin-6 (IL-6), granulocyte colony-stimulating factor (G-CSF), C-X-C motif chemokine ligand 1 (CXCL1), and tumor necrosis factor-alpha (TNFα). These cytokines are well-established mediators of the innate immune response. While no significant differences were found for G-CSF, CXCL1, and TNFα between the two injury models, IL-6 levels were significantly elevated in the C3HC group compared to the C2HS (27.3 ± 4.2 pg/mL vs. 3.2 ± 2.5 pg/mL, p < 0.05), indicating a delayed resolution of systemic inflammation in the contusion model.

## 4. Discussion

In this study, we compared histological, cellular, and inflammatory responses to SCI in two different models of cervical injuries at 7 days post-injury: one by section above the PMN pool (C2HS) and one by contusion at the level of PMN pools (C3HC). Our analysis revealed key differences in neuronal survival, glial activation, and immune cell infiltration that provide insight into the distinct pathophysiological mechanisms triggered by these injuries. C2HS shows limited CSPG accumulation and neuronal preservation, and induces a beneficial profile characterized by Microglial activation, with upregulation of CD86 and F4/80 likely oriented toward repair(Butenko et al., 2020; Sheng et al., 2021; Govindappa and Elfar, 2022; Liu et al., 2022; Zhang et al., 2023) associated with less pro inflammatory cell infiltration compared to C3HC. In contrast, C3HC triggering massive CSPG production, extensive infiltration of immune cells, including Neutrophils and recruited cells exhibiting a dendritic cell–like phenotype. In this context, Microglia remain in a non-homeostatic state (low Tmem119) and do not show marked activation, as CD86 and F4/80 levels remain similar to control levels, suggesting inefficient engagement in microenvironment repair.

These differential immune responses likely reflect variations in the underlying injury mechanisms and could have important implications for targeted therapeutic interventions.

### 4.1 Structural damage and local glial responses following cervical SCI

We found significant distinct alterations in Microglial marker expression between C3HC and C2HS. In the acute phase after SCI, Microglia have a positive impact by clearing debris, recruiting peripheral immune cells into the lesion and releasing cytokines that promote repair (Davalos et al., 2005; Von Bernhardi et al., 2010; Jin and Yamashita, 2016). However, when their activation becomes excessive or prolonged, Microglia fuel a cycle of inflammation that undermines neuronal survival (Fu et al., 2020; Pottorf et al., 2022). Interestingly, while flow cytometry revealed that the number of Microglia at the lesion site was comparable between the two models, the more pronounced Iba+ Microglial response observed below the lesion in C3HC suggests that this model induces a more extensive and widespread neuroinflammatory response compared to C2HS. This increase was accompanied by a significant decrease in Tmem119 expression, a marker associated with homeostatic, non-activated Microglia (Bohnert et al., 2020). This inverse relationship may reflect a shift toward a non-homeostatic, potentially neurotoxic state in the C3HC model (Bohnert et al., 2020; Kenkhuis et al., 2022). In contrast, higher Tmem119 expression in C2HS suggests a more preserved homeostatic Microglial phenotype. These findings suggest that Microglia in the contusion model not only become more broadly activated but may also remain in a prolonged, non-resolving inflammatory state (Zrzavy et al., 2020; Mercurio et al., 2022). While many studies describe Microglial activation as either protective or harmful depending on timing and context, its interpretation *in vivo* should be approached with caution, here we suggest that the nature (contusion vs. section) of the injury plays a key role in shaping their phenotype. Of note, at the lesion site, Microglia from the C2HS model exhibited higher levels of CD86 and F4/80 compared to contusion model. While CD86 is classically considered an M1-associated marker, its upregulation has also been observed in alternatively activated (M2a) phenotype, where increased labeling intensity may indicate comparable antigen density and functional activity, including immune modulation and repair (Liu et al., 2022; Zhang et al., 2023). Similarly, F4/80 is generally expressed at low levels in homeostatic Microglia but can be upregulated during the resolution phase of inflammation, where it is linked to phagocytic capacity and tissue repair (Sheng et al., 2021; Govindappa and Elfar, 2022). The concurrent upregulation of both markers in the C2HS model suggests a transitional state indicative of a repair Microglia-dominated response, potentially reflecting ongoing clearance of debris, an anti-inflammatory or reparative state and resolution of inflammation. This suggests that differences in injury type can strongly influence the balance between resident Microglial activation and peripheral macrophage recruitment, which may have important implications for downstream inflammatory cascades and tissue remodeling. Rather than simply scaling with the lesion size, these immune responses appear to reflect deeper differences in how each injury model perturbs the spinal microenvironment. Our findings emphasize the need to consider the three-dimensional spread and qualitative nature of inflammation when interpreting or targeting immune responses after spinal cord injury. This complex immune landscape not only involves Microglia and infiltrating immune cells, but also engages other glial populations.

Astrocytes closely follow Microglia in the injury cascade. Activated astrocytes proliferate and secrete extracellular matrix molecules, notably chondroitin sulfate proteoglycans (CSPGs), which form perineuronal nets (PNNs) and glial scars (Wiese et al., 2012; Chelyshev et al., 2022). Initial inflammatory cues drive astrocyte activation and CSPG overproduction. Accumulated CSPGs then bind receptors such as PTPσ and toll-like receptor 4 on astrocytes (Ohtake et al., 2016b; Francos-Quijorna et al., 2022), reinforcing GFAP expression and presents a major barrier to post-traumatic neuroplasticity by restricting axonal regrowth or sprouting (John et al., 2023). This feedback loop not only blocks axonal sprouting by interfering with integrin signaling and collapsing growth cones (Bradbury et al., 2002) but also compounds neuronal loss and limits recovery potential (Zhao et al., 2013). In our contusion model, elevated CSPG production coincided with pronounced ventral neuron depletion and spread below the lesion, creating an environment that is hostile to circuit remodeling and likely underlies the poorer functional recovery. By contrast, hemisection spares the integrity of phrenic motoneurons (only a deafferentation on one side) below the lesion level since they are not directly damaged (anatomical distribution at C3-C5) (Gauthier et al., 2006; Michel-Flutot et al., 2021). The combination of neuron depletion and CSPG-rich matrix in C3HC likely impedes circuit remodeling: CSPGs not only form physical barriers but also inhibit integrin-mediated signaling necessary for growth cone advance (Pizzorusso et al., 2002). While our data link CSPG accumulation to more severe pathology, it remains unclear whether CSPGs drive Microglial/astrocytic activation or simply reflect tissue damage. Future studies should employ enzymatic CSPG degradation (e.g., chondroitinase ABC) (Bradbury et al., 2002; Burnside et al., 2018; Jevans et al., 2021) or receptor blockade (PTPσ inhibitors) (Lang et al., 2015) to determine causality and assess functional recovery in both models. Additionally, longitudinal analyses beyond seven days will clarify how CSPG dynamics and glial phenotypes evolve over time and influence chronic remodeling.

### 4.2. Divergent innate Immune responses and systemic inflammation following distinct types of cSCI

SCI is not only a mechanical insult but is also rapidly interpreted by the immune system as a serious biological threat, triggering an innate immune response (Sterner and Sterner, 2023). Our flow cytometry data reveal that, C3HC and C2HS injuries elicit distinct immune profiles, with the C3HC model inducing a more sustained and stronger innate immune response characterized by increased Neutrophils, natural killer (NK) cells, and infiltrating antigen-presenting cells (APCs). This pattern suggests that contusion injuries maintain or amplify inflammatory activity compared to sections: likely due to more severe vascular disruption and secondary tissue damage. In particular, vascular rupture following contusion may lead to local ischemia and reperfusion events, reducing oxygen and glucose delivery and resulting in pronounced metabolic stress in the spinal cord parenchyma. These microenvironmental changes not only promote secondary degeneration but may also act as potent inflammatory triggers that amplify immune cell recruitment (Hu et al., 2023; Li et al., 2023). Such a hostile metabolic milieu might contribute directly to neuronal loss, as observed in our model, or indirectly by sustaining inflammatory cascades. For example, hemisection injuries cause immediate disruption of the blood-spinal cord barrier and rapid immune cell infiltration, whereas contusion injuries often lead to prolonged ischemia, edema, and sustained inflammatory signaling (Clifford et al., 2023). These distinctions may have important implications for therapies aimed at modulating inflammation. The surge in Neutrophils observed in the C3HC group likely reflects an early-stage inflammatory response aimed at debris clearance and tissue repair(Peiseler and Kubes, 2019; Sas et al., 2020), but sustained neutrophil activity may also worsen tissue damage through excessive production of reactive oxygen species and proteolytic enzymes (Dolma and Kumar, 2021). The elevated NK cell count further indicates intensified innate immune activation. NK cells are capable of eliminating compromised cells and modulating immune cascades, playing both protective and pathogenic roles in the CNS (Garofalo et al., 2020; Ning et al., 2023). Their enhanced recruitment in C3HC implies greater need for cytotoxic surveillance and immune regulation under extensive parenchymal stress. Among the infiltrating immune cells, their overall abundance was greater in the C3HC group than in C2HS, suggesting a more intense recruitment of peripheral immune components following contusion. Within this context, we observed a marked increase in CD8⁺ DC-like cells in both injury models compared to controls, which may reflect a general inflammatory adaptation following SCI rather than classical antigen presentation, since our model does not involve an infectious context (Gillespie et al., 2024). In the C3HC group, the elevated expression of CD11b+ DC like cells associated with inflammatory activation (Hu et al., 2024). This phenotype more likely indicates an intensified and potentially harmful inflammatory state at the contusion site, which could contribute to secondary tissue damage and impair repair. Systemic cytokine measurements further support this interpretation. We found significantly elevated systemic IL-6 levels in the C3HC group, while IL-6 levels in C2HS animals were similar to those of controls. They are rapidly upregulated following tissue injury or infection, and contribute to the recruitment and activation of immune cells such as Neutrophils and macrophages, and enhance inflammatory signaling at both the local and systemic levels(Florentin et al., 2021; Soliman and Barreda, 2022; Guan et al., 2025), its persistent elevation at 7 days post-injury in C3HC animals suggests a prolonged inflammatory response. This is in line with previous studies showing that increased IL-6 correlated with more severe injury and reduced neurological recovery (Stammers et al., 2012; Kwiecien et al., 2020). These findings reinforce the notion that contusion injuries lead to sustained peripheral inflammation, in contrast to C2HS, where the immune environment appears to resolve more rapidly and shift toward repair. Consistent with these systemic findings, our immunofluorescence data further demonstrates a more deleterious tissue response in the C3HC model. The accumulation of CSPG, an established inhibitor of axonal regeneration, was markedly higher in C3HC animals, and fewer ventral neurons were preserved. In contrast, the C2HS model showed lower CSPG levels, greater neuronal preservation, and molecular markers suggestive of a reparative macrophage response. Altogether, our data suggests that contusion injuries provoke a more harmful neuroinflammatory and systemic immune response compared to hemisections, which may partially explain the differing regenerative and functional outcomes observed between the models. Moving forward, mechanistic studies using in vivo immune cell tracking, conditional knockout models targeting dendritic-like cell function or IL-6 signaling, and single-cell RNA sequencing of both central and peripheral immune compartments will be essential to dissect the precise cellular crosstalk and molecular pathways driving divergent immune responses after different SCI types.

### 4.3 Clinical Relevance

The differences found between our two models may explain some discrepancy between promising treatment in preclinical models and human reality. For example, anti-Nogo-A therapeutic has been shown to promote corticospinal tract regeneration and functional recovery in hemisection models of cSCI, where axonal growth inhibition represents a major obstacle (Merkler et al., 2001; Sartori et al., 2020). However, this strategy has shown limited benefits in clinical trials, potentially due to the more diffuse damage and multifactorial inhibition seen in human cervical injuries, conditions better reflected in contusion models like hemicontusion. Together, these factors underscore the importance of using appropriate injury models that better represent human pathology when evaluating potential treatments. The combination of increased expression of glial markers (IBA1 and GFAP), sustained systemic inflammation, and elevated CSPG after C3HC adds to the direct damage to the phrenic motor pool after and further compromise respiratory circuitry than C2HS model, in which ipsilateral neural pathways are preserved below the lesion level. Previous studies have linked elevated inflammatory cytokines and glial activation with impaired respiratory outcomes following cervical SCI(Truflandier et al., 2018; Bajjig et al., 2022). Future work should explore whether modulating the inflammatory environment can mitigate respiratory dysfunction models of contusion that are closer to human pathophysiology.

### 4.4 Limitations and Strengths

One limitation of our study is that the analysis was confined to the subacute phase (7 days post-injury), which may not capture the chronic evolution of neuroinflammatory responses and tissue remodeling. However, in clinical settings, patients with cervical spinal cord injury typically experience a delay before receiving rehabilitation. From a mechanistic standpoint, the 7-day timepoint coincides with the resolution phase of initial neutrophil infiltration, the emergence of adaptive immune activity, and the transition toward gliosis and tissue remodeling, critical processes that influence long-term outcomes (Keirstead et al., 2005; Rowland et al., 2008; Hubertus et al., 2025). Nonetheless, characterizing immune responses across multiple time points would provide a more comprehensive understanding of the temporal dynamics underlying spinal cord injury and repair. A second limitation lies in the use of a mouse contusion model; while it effectively replicates many pathophysiological, functional, and morphological features of human SCI, including the absence of spontaneous neural regeneration, it does not exhibit cystic cavity formation, which is typically observed in rat models. This discrepancy may limit direct translational relevance. However, mouse models faithfully replicate key features of human SCI, including the lack of spontaneous regeneration, the presence of both central and peripheral immune activation, and the development of functional motor deficits. Furthermore, mice allow for the use of sophisticated genetic tools and cell-specific tracing techniques that can dissect molecular pathways with high precision. These strengths make the mouse model particularly well-suited for mechanistic investigations that inform potential clinical strategies, even if some anatomical differences limit direct translation (Sharif-Alhoseini et al., 2017).

## 5. Conclusion

In summary, this study aimed to compare two high cervical spinal cord injury models, the well-established C2HS and the clinically relevant C3HC model, in order to gain a clearer understanding of how different injury types shape immune responses and tissue pathology. Cervical SCI presents major clinical challenges due to its impact on both respiratory and motor functions, yet preclinical models often differ in their injury mechanisms and biological outcomes. Notably, our findings reveal that the C2HS model activates Microglia toward a repair function with rapid resolution of systemic inflammation and limited CSPG accumulation at day 7 that spares ventral neurons, whereas the C3HC model directly compromises them, and drives massive CSPG deposition and recruitment of cells adopting a dendritic cell-like phenotype that is less effective at restoring the lesion microenvironment. This profile is accompanied by prolonged systemic inflammation and increased recruitment of inflammatory cells at the lesion site. By highlighting how anatomical targeting, and lesion complexity influence neuroinflammation and neural integrity, our findings provide insight into the divergent secondary injury cascades initiated by sharp versus contusive trauma. These distinctions underscore the need for model-specific therapeutic strategies, especially in the design of interventions targeting neuroinflammation, neuronal protection, and functional recovery. Future studies should build on this work underlying these differences to guide more precise and effective treatments for patients with cervical spinal cord injury.

## CRediT authorship contribution statement

**Wei Chen**: Writing – original draft, Writing – review & editing, Project administration, Investigation, Methodology, Funding acquisition, Formal analysis, Data curation, Conceptualization, Visualization. **Lucille Adam**: Writing – review & editing, Investigation, Project administration, Methodology, Formal analysis, Data curation, Conceptualization, Visualization. **Michel-Flutot Pauline**: Project administration, Investigation. **Arnaud Mansart**: Resources, Funding acquisition. **Stéphane Vinit**: Writing – review & editing, Project administration, Investigation, Resources, Conceptualization, Funding acquisition. **Isabelle Vivodtzev**: Writing – review & editing, Project administration, Investigation, Resources, Funding acquisition, Conceptualization, Supervision.

