## Supplemental Table 1 for "Unveiling Distinct Neuroimmune Responses in Mouse Models of Cervical Spinal Cord Injury: Hemisection versus Hemicontusion"

**SUPPLEMENTARY MATERIAL**

Supplementary Figures

Legends for Supplementary Data

Supplementary Tables

**Supplementary Figure1- Manual gating strategy used for immune cell recruitment assessment.**

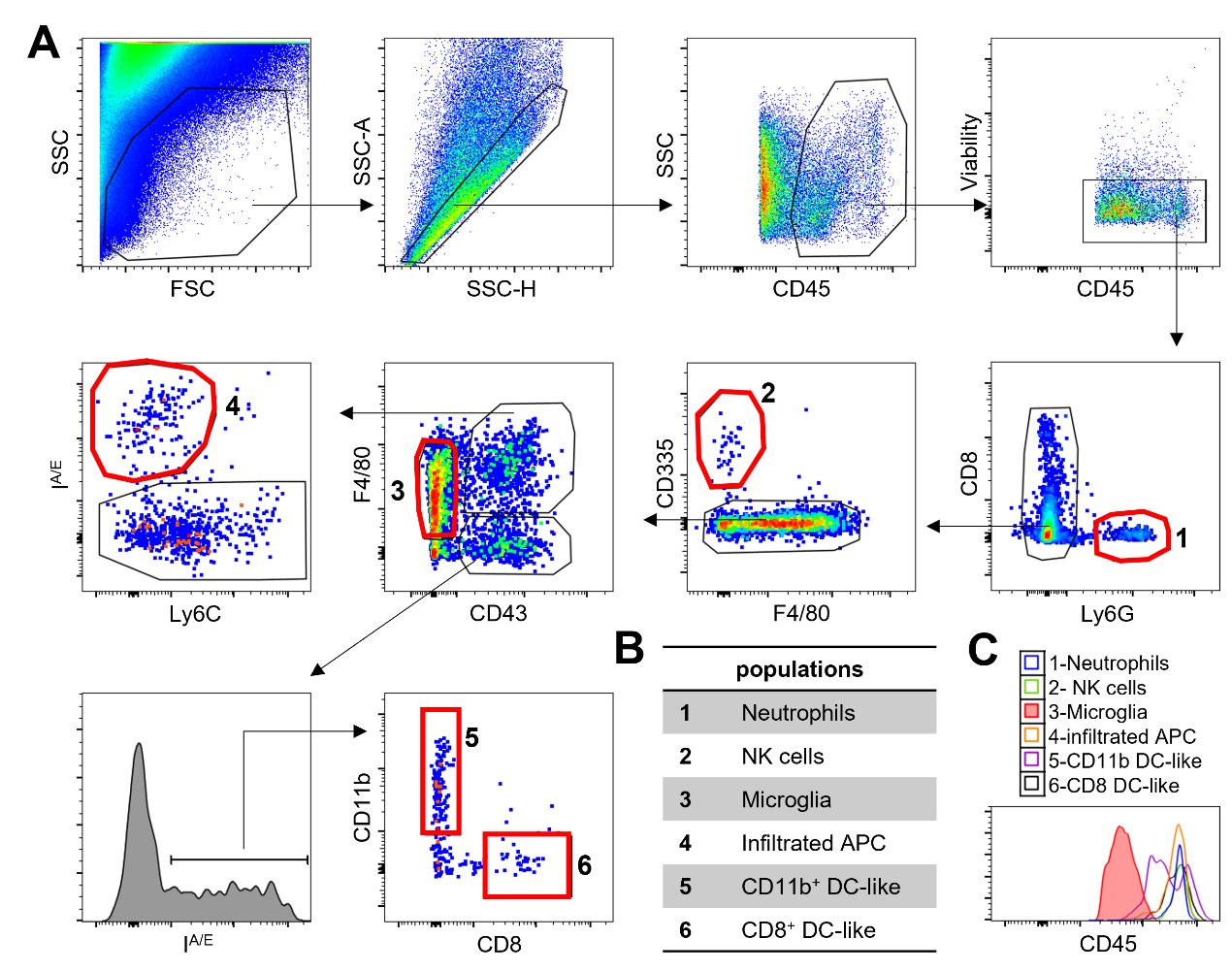

**Supplementary Figure 1**. Characterization of immune cells in spinal cord at lesion or sham site by flow cytometry.

The gating strategy used allow to discriminate immune cells through their CD45 expression, after debris, doublets, and dead cells exclusion. The gating strategy is shown for one representative animal for the hemicontusion condition.

(A). Different immune populations were isolated according to this strategy: neutrophils (Ly6G+), NK cells (CD335+), infiltrating F4/80+ cells and two populations of infiltrating CD43+ F4/80- I^A/E^+ cells, CD11b+CD8- and CD11b+CD8- (A, B). Expression level for CD45 is shown to validate the phenotype of microglia characterized as CD45^low^ (C).

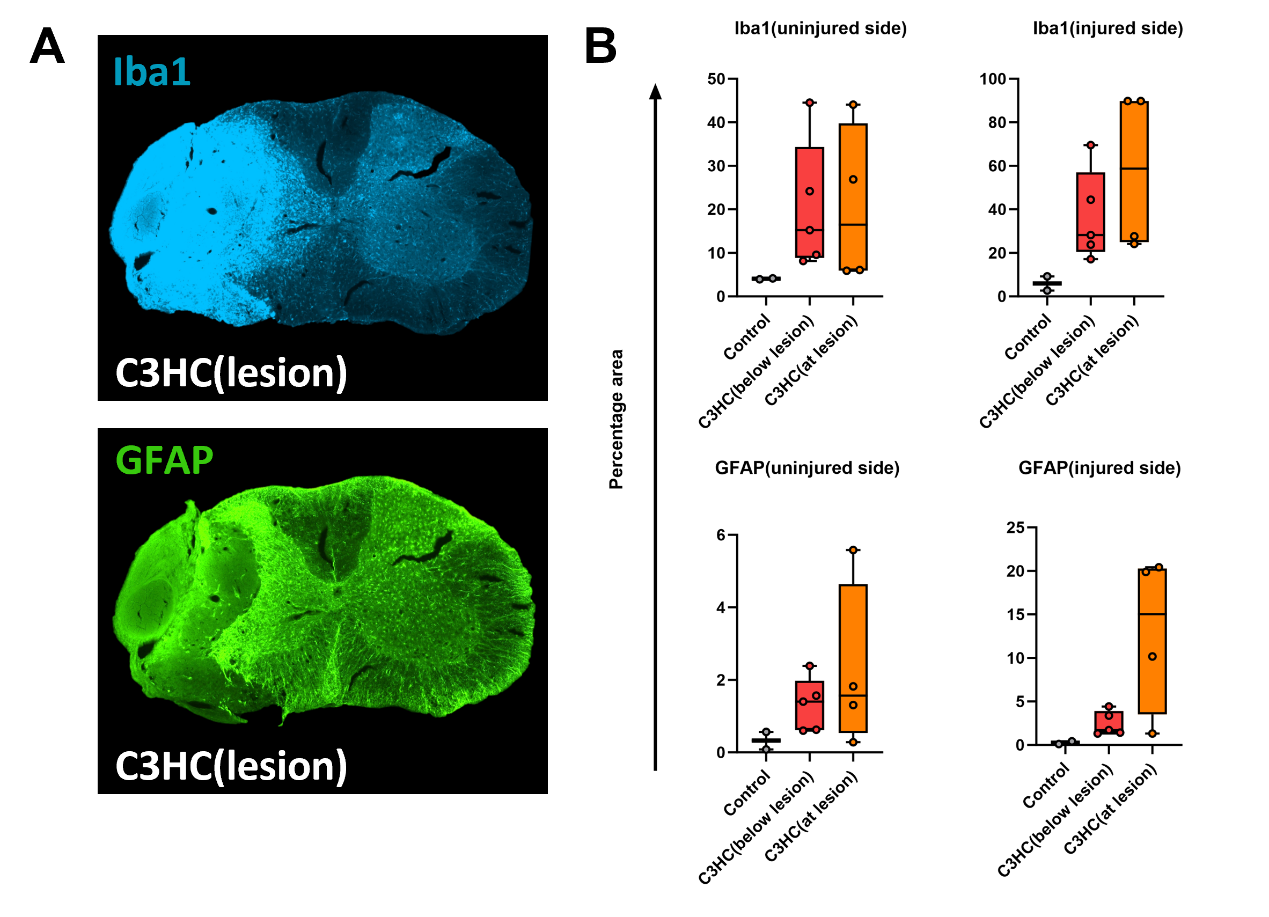

**Supplementary Figure 2**. **Microglial and astroglial activation at 7 days post-injury in C3HC cervical spinal cord injury models at the lesion level.**

**A.** Representative images of expression of ionized calcium-binding adaptor protein-1 (Iba1) (microglia, in blue) positive percentage area at ventral horn and expression of glial fibrillary acidic protein (GFAP) (Astrocyte, in green) positive percentage area at ventral horn for control, C3HC group following 7 days post-injury below and at the lesion level. **B.** Quantification of the percentage positive area of Iba1 and GFAP staining at ventral horn for uninjured and injured sides of C3HC animals following 7 days post-injury at and below the lesion level. Scale bar: 500 µm. Statistical analysis was performed using the One-Way ANOVA and unpaired t test (*), * 𝑝<0.05, ** 𝑝<0.01, *** 𝑝<0.001.

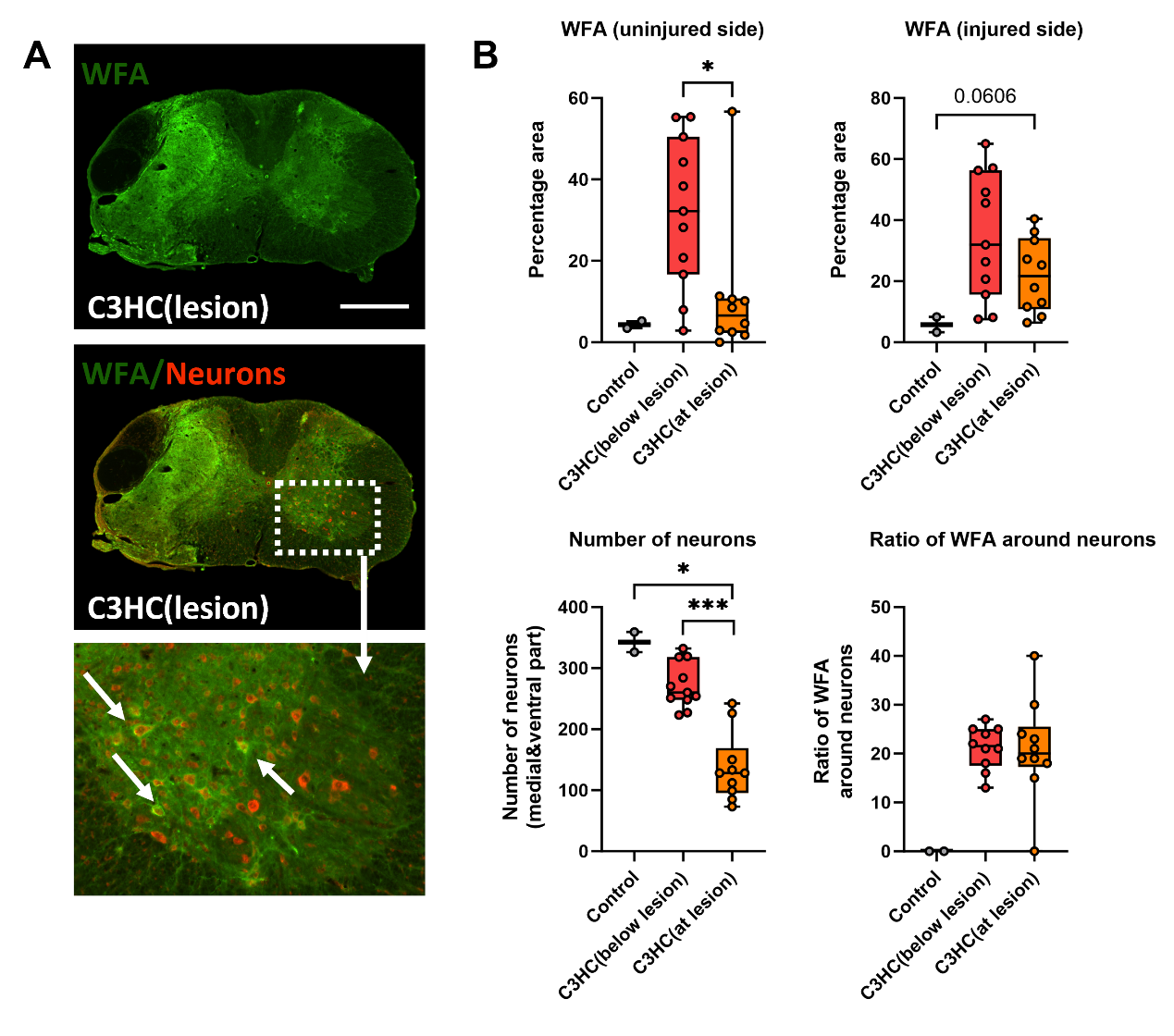

**Supplementary Figure 3**. **CSPG accumulation and ventral neuron changes at 7 days post-injury in C2HS and C3/4HC cervical spinal cord injury models at the lesion level.**

**A.** Representative images of expression of Chondroitin sulfate proteoglycan (CSPG) (wisteria floribunda agglutinin, WFA) and neurons (ventral part) surrounded by WFA for control, C3HC group following 7 days post-injury at the lesion level. **B.** Quantification of the percentage of WFA staining and neurons and ratio of WFA around neurons for uninjured and injured sides of C3HC animals following 7 days post-injury around the lesion level. Scale bar: 500 µm. Statistical analysis was performed using the One-Way ANOVA and unpaired t test (*), * 𝑝<0.05, ** 𝑝<0.01, *** 𝑝<0.001.

**Supplementary Table 1 – Immunochemistry antibodies list**

| **Immunochemistry primary antibodies** | | |
| --- | --- | --- |
| **Antibody** | **Dilution** | **Company - Ref** |
| Goat anti-Iba1 | 1/400 | Abcam ab5076 |
| Rabbit anti- GFAP | 1/4000 | Millipore-Merck AB5804 |
| WFA(Biotinylated wisteria floribunda lectin) | 1/2000 | Vector laboratories |
| **Immunochemistry secondary antibodies** | | |
| **Antibody** | **Dilution** | **Company - Ref** |
| Alexa Fluor 647 Donkey anti-goat | 1/2000 | Molecular Probes A21447 |
| Donkey anti-rabbit Alexa Fluor 488 | 1/2000 | Molecular Probes A21206 |
| Alexa Fluor 488 Avidin | 1/1000 | Molecular Probes A21370 |

**Supplementary Table 2 – Flow cytometry panels**

| **Mix macrophages** | **Dilution** | **Company - Ref ref** |
| --- | --- | --- |
| CD11b PB | 1/800 | Biolegend 101224 |
| IA/IE BV711 | 1/800 | Biolegend 107643 |
| CD45 BV510 | 1/200 | Biolegend 103138 |
| Ly6G AF700 | 1/200 | Biolegend 127617 |
| CD68 AF647 | 1/100 | Biolegend 137004 |
| Ly6C PercPCy5.5 | 1/200 | Biolegend 128012 |
| CD86 FITC | 1/100 | Biolegend 105005 |
| CD206 BV650 | 1/100 | Biolegend 104453 |
| CD43 PE/Dazzle 594 | 1/400 | Biolegend 143217 |
| F4/80 PE | 1/400 | Biolegend 123109 |
| Tmem119 PC7 | 1/100 | Thermofisher |

| **Mix NK cells** | **dilution** | **Company - Ref ref** |
| --- | --- | --- |
| CD11b PB | 1/800 | Biolegend 101224 |
| IA/IE BV711 | 1/800 | Biolegend 107643 |
| CD45 BV510 | 1/200 | Biolegend 103138 |
| Ly6G AF700 | 1/200 | Biolegend 127617 |
| CD335 APC | 1/100 | Biolegend 137608 |
| Ly6C PercPCy5.5 | 1/200 | Biolegend 128012 |
| CD8a AF488 | 1/200 | Biolegend 100726 |
| CD206 BV650 | 1/100 | Biolegend 104453 |
| CD43 PE/Dazzle 594 | 1/400 | Biolegend 143217 |
| F4/80 PE | 1/400 | Biolegend 123109 |
| CD3ε PE/Cyanine7 | 1/100 | Biolegend 155621 |
